# A comparative study of the effects of Aducanumab and scanning ultrasound on amyloid plaques and behavior in the APP23 mouse model of Alzheimer disease

**DOI:** 10.1101/2021.01.04.425327

**Authors:** Gerhard Leinenga, Wee Kiat Koh, Jürgen Götz

**Affiliations:** Clem Jones Centre for Ageing Dementia Research, Queensland Brain Institute, The University of Queensland, Brisbane, QLD 4072, Australia

**Keywords:** Alzheimer’s disease, blood–brain barrier, dementia, amyloid-β, immunotherapy

## Abstract

**Background:** Aducanumab is an anti-amyloid-β (Aβ) antibody that achieved reduced amyloid pathology in Alzheimer’s disease (AD) trials, but it is controversial whether it also improved cognition. It has been claimed that this would require a sufficiently high cumulative dose of the antibody in the brain. Therapeutic ultrasound, in contrast, has only begun to be investigated in human AD clinical trials. We have previously shown that scanning ultrasound in combination with intravenously injected microbubbles (SUS), that temporarily and safely opens the blood-brain barrier (BBB), removes amyloid and restores cognition in APP23 mice. It has not been directly tested how the effects of SUS compare to immunotherapy or whether a combination therapy is more effective.

**Methods:** In a study comprising four treatment arms, we tested the efficacy of an Aducanumab analogue, Adu, in comparison to SUS, as well as a combination therapy in APP23 mice, using sham as a control (aged 13-22 months). The active place avoidance (APA) test was used to test spatial memory, and histology and ELISA were used to measure amyloid. Brain antibody levels were also determined.

**Results:** We found that both Adu and SUS reduced the total plaque area in the hippocampus to a similar degree, with no additive effect in the combination treatment (SUS+Adu). Whereas there was only a trend towards a reduction for both Adu and SUS in the cortex, the combination trial yielded a statistically significant reduction compared to sham. Only the SUS and SUS+Adu groups included animals that had their plaque load reduced to below 1% from above 10%. There was a robust improvement in spatial memory for SUS+Adu only. In this group, when measured three days post-treatment, Adu levels were still 5-fold increased in the combination therapy compared to delivery of Adu on its own.

Together, these findings suggest that SUS should be seriously considered as a treatment option for AD. Alternatively, a combination trial using Aducanumab together with ultrasound to increase brain levels of Aducanumab may be warranted, as the two approaches may engage different (albeit shared) clearance mechanisms.

## Background

The deposition of amyloid-β (Aβ) in the brain is considered a key initiating step in the development of Alzheimer’s disease (AD). Approaches that either prevent or remove the accumulation of Aβ in the brain have been a focus of research into developing a therapy for this disorder [1–3]. Several active and passive immunization strategies targeting Aβ have been explored in clinical trials for AD to enhance Aβ clearance from the brain.

Aducanumab is an anti-Aβ antibody that targets Aβ aggregates including insoluble fibrils and soluble oligomers by binding to the amino-terminus of Aβ at residues 3-7 in a shallow pocket in the antibody [4]. This human IgG1 antibody had been isolated from the B cells of cognitively healthy elderly humans and has low affinity for monomeric Aβ [5]. In a Biogen-sponsored phase 1b clinical trial (PRIME) of Aducanumab in prodromal and mild AD patients, a striking reduction in amyloid plaques as measured by positron emission tomography (PET) was reported following one year of monthly intravenous antibody infusions at doses ranging from 3-10 mg/kg. One of the two phase III trials of Aducanumab, EMERGE, unlike ENGAGE, showed reductions in cognitive decline, possibly reflecting the effects of higher accumulated doses of the antibody [6]. Biogen is currently seeking U.S. Food and Drug Administration (FDA) approval for Aducanumab and may be granted conditional approval of the therapy pending a post-market commitment of a phase IIIB re-dosing trial that has recently been launched. If approved, it will be the first anti-amyloid agent and first antibody treatment for AD. However, it remains to be determined whether Aducanumab is a disease-modifying therapy that achieves significant clinical benefits in AD patients [7]. The interpretation of the cognitive data from these trials is complex because dosing was altered and stopped during the trial, and the magnitude of the cognitive effect was relatively small. Sevigny *et al* demonstrated 50% plaque reduction in 9.5-15.5-month-old amyloid precursor protein (APP) mutant Tg2576 mice after treating with a mouse IgG2a Aducanumab analog; however, the effects of the immunotherapy on behavioral read-outs in mice have not been reported.

Therapeutic ultrasound is an alternative strategy for clearing amyloid by transiently opening the blood-brain barrier (BBB) and allowing for the uptake of blood-borne factors and therapeutic agents [8]. Given that ultrasound parameters are highly tunable, this technique can be safely applied to a range of species, including mice [9, 10], dogs [11], sheep [12, 13], macaques [14, 15] and also humans [16]. Even without using a therapeutic agent (such as an antibody), repeated opening of the BBB with the scanning ultrasound (SUS) approach in 12 and 22 month-old APP23 mice has been shown to activate microglia and thereby reduce amyloid and improve memory performance [17, 18], that was dependent on BBB opening rather than simply applying ultrasound as a pressure wave to induce neuromodulatory effects [19]. The underlying mechanisms of ultrasound-mediated BBB opening have not been fully dissected but involve both facilitated para- and transcellular transport [20]. In preclinical studies, ultrasound was used to deliver model molecules of various sizes [21], as well as antibodies [22–24] to the brain.

Here, we sought to compare the efficacy of treatment with an Aducanumab analog (Adu), SUS and a combination of both SUS and Adu regarding plaque burden and performance in a spatial memory task.

## Materials and Methods

### Study design

APP23 mice express human APP751 with the Swedish double mutation (KM670/671NL) under the control of the neuron-specific mThy1.2 promoter. As they age, these mice exhibit memory deficits [25], amyloid plaque formation initiating in the cortex, and cerebral amyloid angiopathy (CAA) [26]. In this study, APP23 mice, aged 13 months, were assigned to four treatment groups: sham (N=10), SUS (N=11), Adu (5 mg/kg delivered retroorbitally, N=11), or SUS+Adu (5 mg/kg retroorbitally, N=10). Assignment to treatment groups was based on matching performance of spatial memory (number of shocks) on day 5 of the active place avoidance (APA) test. A group of wild-type mice (N=12) were included. APP23 mice were ranked from those receiving the fewest shocks to those receiving the most shocks on day 5 and were assigned to the four treatment groups (sham, SUS, Adu, SUS+Adu) in rank order. Each group received a total of nine treatments (an APA retest was performed after the forth treatment), with the final treatment in the Adu and SUS+Adu groups using fluorescently labelled antibody (2.5 mg/kg Alexa Fluor 647-labeled Adu and 2.5 mg/kg unlabeled Adu). (**Fig. 1A**). Three days after the final treatment, the mice were administered an overdose of sodium pentobarbitone and perfused with phosphate buffered saline (PBS). The right hemisphere of the brain was fixed in paraformaldehyde solution for histology, while the cortex and hippocampus of the left hemisphere were dissected and frozen in liquid nitrogen for subsequent analysis. Due to the increased mortality of this strain [27], the numbers of mice surviving to 22 months for histological and biochemical analysis were N=10 sham, N=9 Adu, N=8 SUS, N=9 SUS+Adu. Assessment of outcomes was performed with the researcher blinded to the treatment group. All animal experimentation was approved by the Animal Ethics Committee of the University of Queensland (approval number QBI/554/17). Sample sizes for the experiment were selected based on our earlier studies [17]. Data was collected for all mice that survived until the end of the experiment and all data was included.

**Fig. 1.**
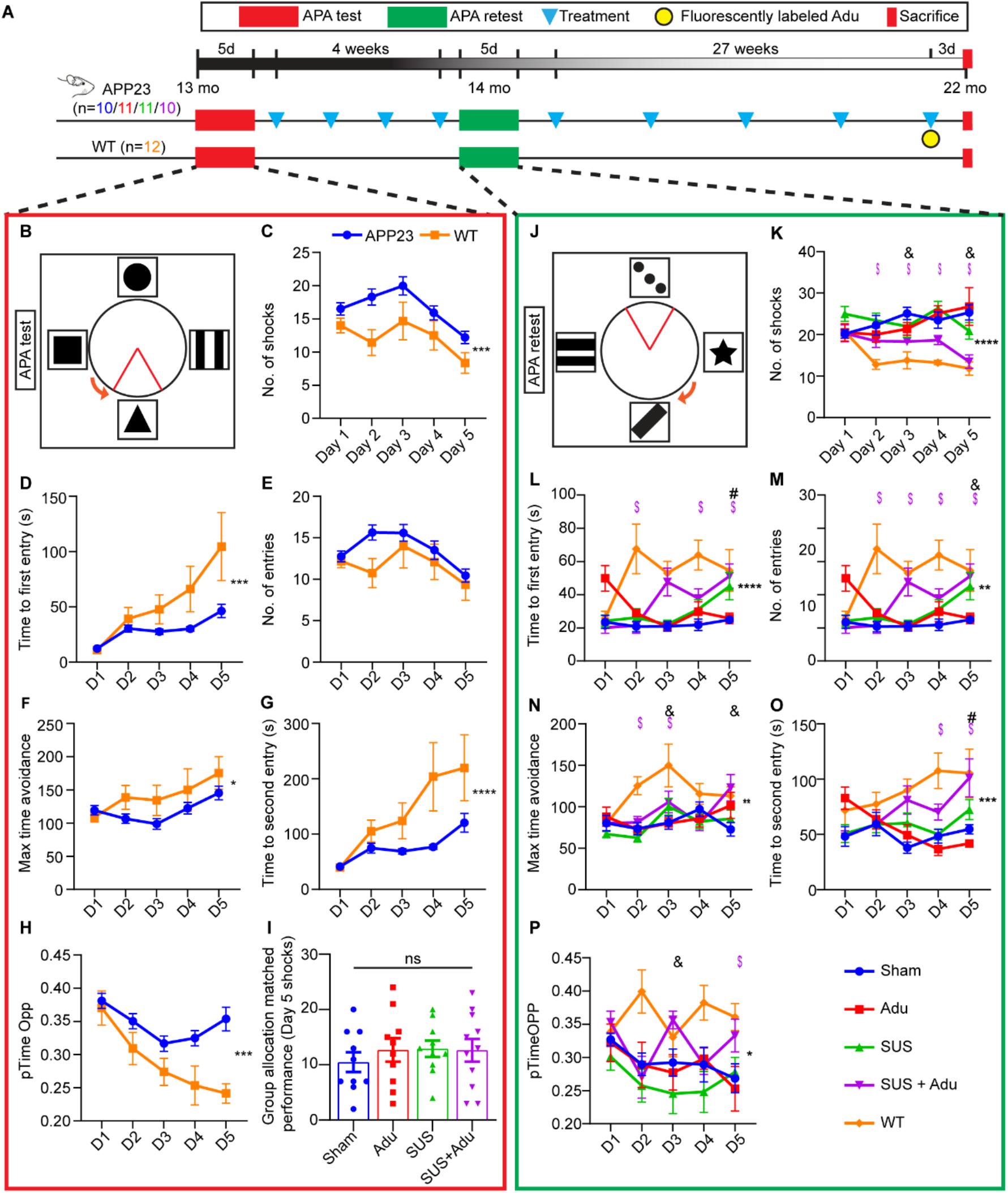
Study overview and results of APA test and retest. (**A**) Overview of the study with timeline. (**B**) Active place avoidance test (APA) in which mice must use spatial cues to avoid a shock zone indicated as a red triangle. (**C**) APP23 mice had impaired performance in the APA test in terms of number of shocks received. (**D**) APP23 mice had shorter time to first entry to the shock zone as determined by two-way ANOVA. (**E**) APP23 mice did not show significant impairment in the measure number of entries (**E**) or maximum time of avoidance (**F**), but were impaired on the measures time to second entry (**G)** and proportion of time spent in the opposite quadrant to the shock zone (**H**). APP23 mice were then assigned to treatment groups based on matched performance on day 5 of the APA (**I**). APA retest was performed after four once-per-week treatments with changes to room cues, shock zone location, and the direction of rotation (**J**). Effect of treatment on number of entries (**M**), maximum time of avoidance (**N**), time to second entry (**O**) and proportion of time spent in the opposite quadrant to the shock zone (**P**). Data is represented as mean ±SEM. Statistical differences: *p<0.05, **p<0.01, ***p<0.001, ****p<0.0001, $=simple effect comparing wild-type vs sham p<0.05, #=simple effect comparing SUS vs sham p<0.05, &=simple effect comparing SUS+Adu vs sham p<0.05. Sham N=10, Adu N=11, SUS N=11, SUS+Adu N=10, WT N=12. Data were analyzed with a two-way ANOVA and follow-up Holm-Sidak tests for simple effects.

### SUS equipment

An integrated focused ultrasound system (Therapy Imaging Probe System, TIPS, Philips Research) was used. This system consisted of an annular array transducer with a natural focus of 80 mm, a radius of curvature of 80 mm, a spherical shell of 80 mm with a central opening of 31 mm diameter, a 3D positioning system, and a programmable motorized system to move the ultrasound focus in the x and y planes to cover the entire brain area [17]. A coupler mounted to the transducer was filled with degassed water and placed on the head of the mouse with ultrasound gel for coupling, to ensure unobstructed propagation of the ultrasound to the brain.

### Production of microbubbles

Microbubbles comprising a phospholipid shell and octafluoropropane gas core were prepared in-house. 1,2-distearoyl-*sn*-glycero-3-phosphocholine (DSPC) and 1,2-distearoyl-*sn*-glycero-3-phosphoethanolamine-N-[amino(polyethylene glycol)-2000] (DSPE-PEG2000) (Avanti Polar Lipids) were mixed in a 9:1 molar ratio, dissolved in chloroform (Sigma) and the chloroform solvent was evaporated under vacuum. The dried phospholipid cake was then dissolved in PBS with 10% glycerol to a concentration of 1 mg lipid/ml and heated to 55°C in a sonicating water bath. The solution was placed in a 1.5 ml glass HPLC vial with the air in the vial replaced with octafluoropropane gas (Arcadophta). The microbubbles were activated on the day of the experiment by agitation of the vial in a dental amalgamator at 4,000 rpm for 45s. Activated microbubbles were measured with a Multisizer 4e coulter counter which reported a mean diameter of 1.885 µm and a concentration of 9.12×10^8^ microbubbles/ml. These microbubbles were also observed to be polydisperse under a microscope (**fig. S2)**.

### SUS application

Mice were anesthetized with ketamine (90 mg/kg) and xylazine (6 mg/kg) and the hair on their head was shaved and depilated. They were then injected retro-orbitally with 1 µl/g body weight of microbubble solution and placed under the ultrasound transducer with the head immobilized. Parameters for the ultrasound delivery were 1 MHz center frequency, 0.7 MPa peak rarefactional pressure, 10 Hz pulse repetition frequency, 10% duty cycle, and a 6 s sonication time per spot. The focus of the transducer was 1.5 mm × 12 mm in the transverse and axial planes, respectively. The motorized positioning system moved the focus of the transducer array in a grid with 1.5 mm spacing between individual sites of sonication so that ultrasound was delivered sequentially to the entire brain as described previously [17, 18]. Mice typically received a total of 24 spots of sonication in a 6×4 raster grid pattern. For the sham treatment, mice received all injections and were placed under the ultrasound transducer, but no ultrasound was emitted. When the animals were treated with Adu antibody, the solution was mixed briefly with the microbubble solution and injected into the retro-orbital sinus before the mouse was placed under the ultrasound transducer. The time between injecting microbubbles and commencing ultrasound delivery was 60 ±10 s and the duration of sonication was approximately 3 min (total time from microbubble injection approximately 4 min).

### Production of the Aducanumab analog

VH and VL sequences were identified in Biogen Idec’s patent submission for BIIB-037 WO2014089500 A1 and were cloned into mouse IgG2a and kappa pcDNA3.1 vectors (GenScript). Murine chimeric Aducanumab (Adu) was produced using the Expi293 expression system, purified using protein A chromatography, and verified to be endotoxin-free by LAL assay (Thermo Fisher).

### Antibody affinity ELISA

The EC_50_ of Adu was determined by direct-binding ELISA. Aβ_1-42_ fibrils were generated by incubating 0.1mM Aβ_1-42_ peptide (JPT Peptide Technologies, Germany) in 10 mM HCl for 3d at 37°C. A MaxiSorp ELISA plate was coated with 2μ/ml Aβ_1-42_ fibrils in 0.1M sodium bicarbonate buffer and then blocked with 1% bovine serum albumin. The EC_50_ was determined by incubating the wells with serial dilutions of Adu, followed by washing and detection of bound Adu with a rabbit anti-mouse horse radish peroxidase conjugated antibody (Dako) and 3,3′,5,5′-tetramethylbenzidine substrate. The 6E10 antibody [28] was used as a positive control for Aβ binding and its EC50 was determined for comparison with Adu using the same methods (**fig. S1)**.

### Antibody labeling

Adu was covalently conjugated with Alexa Fluor 647 dye (Thermo Fisher Scientific) in PBS with 0.1 M sodium bicarbonate as previously described [22]. The protein concentration and degree of labeling were determined by measuring absorbance at 280 nm and 650 nm, respectively.

### Tissue processing

Mice were deeply anesthetized with pentobarbitone before being perfused with 30 ml of PBS, after which brains were dissected. One hemisphere of the brain was fixed overnight in a solution of 4% wt/vol paraformaldehyde, then cryoprotected in 30% sucrose and sectioned coronally at 40 µm thickness on a freezing-sliding microtome (SM2000R, Leica). A one-in-eight series of sections was stored in PBS containing 0.01% sodium azide at 4°C until staining.

### Assessment of amyloid plaques

For the assessment of amyloid plaque load, an entire one-in-eight series of coronal brain sections of one hemisphere at 40 μm thickness was stained using the Campbell-Switzer silver stain protocol that discriminates fibrillar from less aggregated amyloid as previously described [17]. Stained sections were mounted onto microscope slides and imaged with a 10x objective on a Metafer bright-field VSlide scanner (MetaSystems) using Zeiss Axio Imager Z2. Analysis of amyloid plaque load was performed on all stained sections using ImageJ. Separate regions-of-interest were drawn around the cortex and dorsal hippocampus. As both black and amber plaques are present in the sections and represent different types of amyloid compactness, they were analyzed separately using color deconvolution method and automated thresholding to distinguish the two types of amyloid plaques. For the analysis of black plaques, a color deconvolution vector where R=0.65, G=0.70, B= 0.29 was used followed by the MaxEntropy auto thresholding function in ImageJ. As black plaques consist mainly of diffuse fibrils no size filter was applied. To measure amber plaques, a color deconvolution vector where R=0.64, G=0.72 and B=0.27 was used, followed by invert function and automated thresholding using the triangle method in ImageJ, fill-holes function and a 60 μm^2^ size filter was applied. Using this method, plaque number, total plaque area, average plaque size, and % area covered by plaque were obtained for both the black and amber plaques and summed to give total plaque area for the cortex and hippocampus. We were unable to analyze the hippocampus of one mouse in Adu-treated group because of folds in the tissue.

### Assessment of cerebral amyloid angiopathy

To assess CAA, a one-in-eight series of Campbell-Switzer silver-stained sections were examined. Regions of interest were drawn manually around areas of CAA in the cortex, which were distinguished from plaques by having a rod-like structure indicative of blood vessels and a diameter greater than 15 μm. Meningeal CAA which has a ring-shaped structure and occurred close to the edge of the section was also measured. The number of CAA deposits per section, the average size and the % area of brain section positive for CAA staining were determined.

### Immunofluorescence

Coronal 40 μm sections were co-stained with the 4G8 antibody against Aβ (1:1,000, Covance) and Iba1 (1:1,000 Wako) followed by goat anti-mouse and goat anti-rabbit Alexa Fluor-conjugated secondary antibodies (1:2000, Thermo Fisher). Alexa Fluor 647-conjugated Adu was detected in situ without additional amplification. Sections were cover-slipped and imaged with a fluorescence slide scanner (Metafer).

### Enzyme-linked immunosorbent assay for Aβ

Frozen cortices were homogenized in 10 volumes of a solution containing 50 mM NaCl, 0.2% diethylamine (DEA) with complete protease inhibitors, and dounce homogenised by passing through 19 and 27 gauge needles. Samples were then centrifuged at 21,000 × g for 90 min at 4°C. The supernatant was retained as the DEA-extracted soluble Aβ fraction. The remaining pellets were resuspended in 10 volumes of 5 M Guanidine HCl, sonicated, and centrifuged at 21,000 × g for 30 min at 4°C. The resultant supernatant was retained as the guanidine-extracted insoluble Aβ fraction. The concentrations of Aβ_40_ and Aβ_42_ were determined in brain lysates using the ELISA kits according to the manufacturer’s instructions (human Aβ_40_ and Aβ_42_ brain ELISA, Merck).

### Active place avoidance test

The active place avoidance (APA) task is a test of hippocampus-dependent spatial learning. APP23 mice and non-transgenic littermate controls were tested over six days in a rotating elevated arena (Bio-Signal group) that had a grid floor and a 32 cm high clear plastic circular fence enclosing a total diameter of 77 cm. High-contrast visual cues were present on the walls of the testing room. The arena and floor were rotated at a speed of 0.75 rpm, with a mild shock (500 ms, 60 Hz, 0.5 mA) being delivered through the grid floor each time the animal entered a 60-degree shock zone, and then every 1,500 ms until the animal has left the shock zone. The shock zone was maintained at a constant position in relation to the room. Recorded tracks were analyzed with Track Analysis software (Bio-Signal group). A habituation session was performed 24 h before the first training session during which the animals were allowed to explore the rotating arena for 5 min without receiving any shocks. After the first training session, APP23 mice were divided into four groups with mice matched so that the performance (number of shocks) of the four groups of mice on day 5 of the task was the same. A total of five training sessions were held on consecutive days, one per day with a duration of 10 min. Following four weekly treatments, the mice underwent the APA test again (reversal learning). The retest was held in the same room as the initial test. However, the shock zone was switched to the opposite side of the arena, and the visual cues were replaced with different ones, and the platform was rotated clockwise rather than counterclockwise. The number of shocks, numbers of entries, time to first entry, time to second entry, proportion of time spent in the opposite quadrant of the shock zone for sham, SUS, Adu, and SUS+Adu-treated groups were compared over the days of testing.

### Statistical analysis

Statistical analyses were conducted with Prism 8 software (GraphPad). Values were always reported as means ± SEM. One-way ANOVA followed by the Holm-Sidak multiple comparisons test, or t-test was used for all comparisons except APA analyses where two-way ANOVA with day as a repeated measures factor and group as a between subjects factor was performed and was followed by the Holm-Sidak multiple comparisons test for simple effects to compare group performance on different days. The model assumption of equal variances was tested by Brown-Forsyth or Bartlett tests, and the assumption of normality was tested by Kolmgorov-Smirnov tests and by inspecting residuals with QQ plots. All observations were independent, with allocation to groups based on active place avoidance where mice were ranked on performance and assigned to one of the four groups (sham, sus, adu, adu+sus) in order of number of shocks on day 5 listed from most to least shocks.

## Results

### Generation of Aducanumab analog and application

The Aducanumab analog Adu was generated by grafting the VH and VL chains of Aducanumab onto a mouse IgG backbone and expression in Expi293 cells. We then established that the affinity of Adu to fibrillar Aβ_42_ (EC_50_ 81.7 pM) was similar to that published earlier for Aducanumab (EC_50_ 100 pM) [5]. In comparison, the 6E10 antibody had an EC50 of 1.18 nM for Aβ_42_ fibrils (**fig. S1)**. Next, 13-month old APP23 mice were divided into four groups (Adu/SUS/SUS+Adu/sham) based on matching performance on day 5 (final day) of the APA test. A dose of 5 mg/kg Adu was given for each treatment, except for the last treatment where a mixture of 2.5 mg/kg unlabeled Adu and 2.5 mg/kg Alexa Fluor 647-labeled Adu was administered. The mice were initially treated once a week for four weeks, after which they were re-tested in the APA. From 15-22 months of age the mice were subsequently treated five times, then sacrificed three days following the last treatment, resulting in a total of nine treatments (**Fig. 1A**).

### Aducanumab analog when delivered by SUS improves spatial memory performance

In the current study, we compared the effect of delivering the murine chimeric IgG2a Aducanumab analog, Adu, with a SUS treatment, using plaque burden and behavior as the major read-outs. We also assessed a combination treatment (SUS+Adu). Additional comparisons were made by including sham-treated mice, as well as untreated wild-type littermate controls.

We first tested 13 month-old APP23 mice and their wild-type littermates in the APA test of hippocampus-dependent spatial learning in which the animals must use visual cues to learn to avoid a shock zone located in a rotating arena (**Fig. 1B**). Spatial learning could have been assessed in the Morris water maze; however, this test is stressful to mice, and aged mice are poor swimmers [29, 30]. To determine the effect of each treatment protocol on spatial memory function, an APA test consisting of 5 training days with a single 10 min training session each day was performed following habituation to the arena in a one 5 min session the day before the first training day. A two-way ANOVA based on the number of shocks that were received revealed a significant effect of day of testing, indicating that learning had occurred (*F*_4,208_) = 5.728, p= 0.0003. There was also a significant effect of genotype, with APP23 mice receiving more shocks than their wild-type littermates (*F*_1, 52_) = 6.278, p=0.0154 (**Fig. 1C**). There was no significant interaction. Similarly, based on the measure of time to first entry of the shock zone, there was a significant effect of day, with mice showing longer latencies to the first entrance as the number of training days increased (F_4,208_) = 7.586, p=0.0007. Wild-type mice showed longer latencies to enter the shock zone over the days of testing and there was a significant effect of genotype on time to first entry (*F*_*1.52*_) *=* 5.950, p=0.0182 (**Fig. 1D**). Wild-type and APP23 mice did not differ; however, on number of entries (**Fig. 1E**) or maximum time of avoidance (**Fig. 1F**). APP23 mice performed significantly worse on the measures ‘time to second entry’ (**Fig. 1G**) and ‘proportion of time spent in the quadrant opposite to the shocked quadrant’ (**Fig. 1H**) The APA performance of the APP23 mice varied significantly so they were assigned to each of the four treatment groups based on matching performance in terms of the number of shocks received on day 5 of the APA to reduce any differences in performance between treatment groups so as to more readily detect any improvement caused by a treatment (**Fig. 1I**).

Before retesting in the APA, mice were subsequently treated once a week for four weeks. For the retest, the shock zone was shifted by 180 degrees, the cues in the room were changed and the arena rotated in the opposite direction. To perform well in the retest, the mice needed to update their spatial learning in order to learn the new shock zone location (**Fig. 1F**). A two-way ANOVA with group as a between-subjects factor and day as a repeated measures factor revealed a significant effect of treatment group on number of shocks received (*F*_4,47_ = 8.5, p<0.0001). Follow-up multiple comparisons tests showed that SUS+Adu-treated mice received significantly fewer shocks than sham-treated control mice on days 3 (p=0.0295) and 5 (p=0.0005) (**Fig. 1K**). Comparing just the Adu-treated to the SUS+Adu-treated mice by two-way ANOVA revealed that the combined treatment of SUS+Aducanumab led to significantly improved performance over Aducanumab only in terms of number of shocks received during the test (*F*_1,17_ = 6.23, p=0.0231). A two-way ANOVA revealed a significant effect of group (*F*_4,47_ = 6.8, p=0.0002) on time to first entry into the shock zone, with follow-up multiple comparisons test showing that SUS+Adu-treated mice had a longer latency to enter the shock zone on day 5 of the APA task (p=0.024) (**Fig. 1L**). SUS+Adu-treated mice also performed significantly better than sham-treated mice on the measures number of entries (**Fig. 1M**) maximum time of avoidance (**Fig. 1N**), time to second entry (**Fig. 1O**) and proportion of time spent in the quadrant opposite the shock zone (**Fig. 1P**). These results demonstrate that APP23 mice exhibit an improvement in spatial memory when treated with a combination of SUS and Adu. Of note, we wanted to determine amyloid pathology at an advanced age; where it is not possible to perform a third APA test as mice of this age are in a poor condition preventing them from physically performing the task.

### Comparison of plaque reduction in the cortex for the different treatment groups

Following APA testing to ascertain the effects of four once-per-week treatments on spatial memory performance, APP23 mice had five treatment sessions between the age of 15 and 22 months in order to determine whether treating them with SUS, Adu alone, or a combination resulted in robust plaque removal, even at older ages when plaque burden is maximal. The mice were sacrificed at 22 months of age, three days after the last treatment and one hemisphere was processed for histology to identify plaques. We performed Campbell-Switzer staining, which can differentiate diffuse and compact species of amyloid plaques in the brain and is not confounded by the binding of Adu to Aβ. This revealed a reduction in the total amount of plaque area stained when comparing the treatment groups to sham controls **(Fig. 2A).** We calculated the percentage area occupied by plaque for two regions of interest, the cortex and the hippocampus, in 15-20 sections per mouse, assessing plaque burden in a one-in-eight series of sections throughout the rostral-caudal extent of the brain. Analysis of cortical plaque burden in the different groups revealed an effect of treatment (F_3,31_=3.78, p=0.02). A follow-up Holm-Sidak test found that combined SUS+Adu treatment resulted in a statistically significant 52% plaque reduction in the cortex of SUS+Adu-treated APP23 mice compared to sham (p=0.0066). At 22 months of age, these mice have a severe plaque burden, with diffuse and compact plaques occupying 23% of the cortex in the sham-treated mice, compared to 16% of the cortex in mice administered Adu, 17% in mice administered SUS only and 11% in mice which received a combination SUS+Adu treatment (**Fig. 2B**). As an additional analysis, the SUS+Adu treatment was found to be superior to Adu alone in reducing plaque burden (one-tailed t-test, p=0.029). As the Campbell-Switzer silver differentiates diffuse plaques stained black with a cotton wool appearance, and compact plaques stained amber (**Fig. 2A**), we also performed an analysis of these plaques separately using a color deconvolution method in ImageJ. The results of this analysis revealed that the reduction in total plaque area was largely driven by a reduction in the total area of black plaque which occupied 18.60% of the cortical area in sham-treated mice compared to 7.93% in SUS+Adu-treated mice (p=0.0119) (**Fig. 2C**). In contrast, we found no significant difference between the treatment groups based on the area, number, or size of amber plaques (**Fig. 2D,E,I**).

**Fig. 2.**
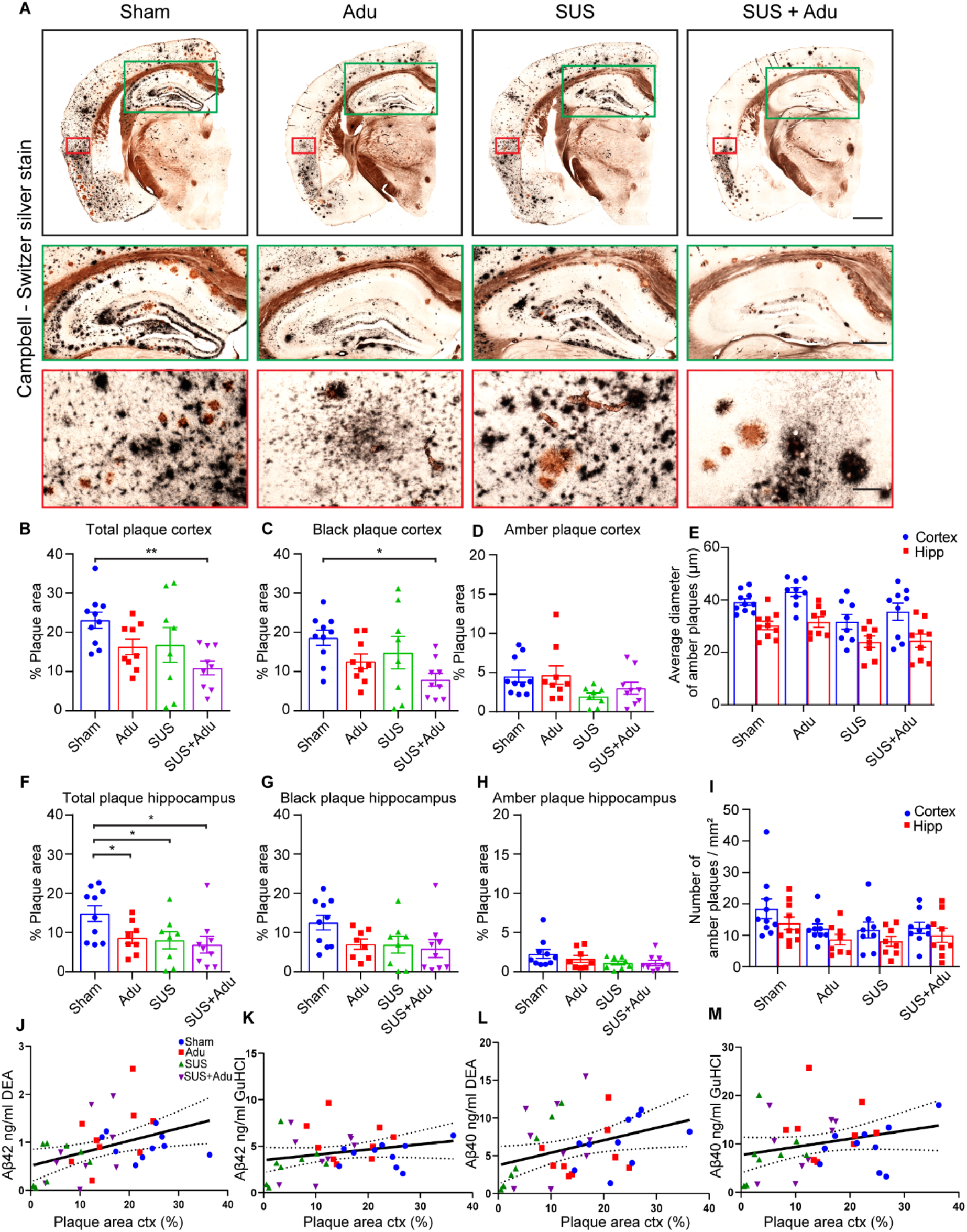
Treatment strategies reduce plaques in APP23 mice. (**A**) Representative Campbell-Switzer silver staining for amyloid plaques in the four treatment groups. Plaques stained black are more diffuse, while amber plaques are compact and discrete. The black box shows the entire hemisphere (scale bars: 1mm). Insert outlined in green shows higher magnification view of dorsal hippocampus (scale bars: 500 μm), and the red inset shows higher magnification image of temporal cortex (scale bars: 100 μm). (**B**) There was a significant reduction of plaques in the cortex of SUS+Adu-treated mice, driven largely by reduction in black plaques (**C**) as area, number and size of amber plaque was less affected by treatment (**D, E,I**). Plaque load in the hippocampus was reduced by Adu, SUS and SUS+Adu (**F**) and hippocampal black plaques (**G**) and amber plaques (**H**) were analyzed separately. A significant correlation was found between amyloid levels measured by histology and by ELISA (**J-M**). Data is represented as mean ±SEM. Statistical differences: *p<0.05, **p<0.01, ***p<0.001. Data were analyzed with a one-way ANOVA and follow-up Holm-Sidak tests. Sham N=10, Adu N=9, SUS N=8, SUS+Adu N=9.

### Aducanumab analog, SUS and the combination therapy all effectively reduce amyloid plaques in the hippocampus of APP23 mice

In APP23 mice, plaque formation is initiated in the cortex and then proceeds to the hippocampus. When we analyzed the total plaque burden in the hippocampus, we found a significant effect of treatment (F_3,31_=, p=0.03, one-way ANOVA). All three treatments (Adu, SUS and Adu+SUS) led to a significant reduction in total plaque area in the hippocampus compared to that in sham-treated APP23 mice (**Fig. 2F)**. The sham-treated mice had a hippocampal plaque burden of 14.84% vs 8.68% (p=0.0432) for Adu-treated mice, 8.04% for SUS-treated mice (p=0.043) and 6.92 % for SUS+Adu-treated mice (p=0.022). However, unlike the effects seen in the cortex, the reduction in total plaque in the hippocampus was not disproportionately driven by a reduction in black plaque (*F*_3,30_=2.43, p=0.08) (**Fig. 2G**) compared to amber plaque reduction (*F*_3,30_ =1.80 p=0.17) (**Fig. 2H**) as it was only when total plaque burden was analyzed that statistically significant reductions in plaques were found. These results show that the effect of SUS on reducing plaque burden in the hippocampus is comparable to that of Aducanumab. Delivering both Aducanumab and SUS together did not significantly improve the plaque clearing ability in the hippocampus of the treatments by themselves.

### Effects of treatment on amyloid-β species

We further performed ELISA measurements of Amyloid-β40 and Amyloid-β42 species from the lysate from one cortex, fractionating proteins into a DEA fraction containing soluble proteins and a guanidine fraction containing insoluble proteins. Levels of Aβ42 and Aβ40 in the DEA soluble fraction were significantly correlated to plaque burden as measured by Campbell-Switzer silver staining of the left hemisphere (R^2^=0.17, p=0.014 and R^2^=0.13, p=0.030 respectively) (**Fig. 2J,L**). Levels of Aβ42 and Aβ40 in the guanidine fraction containing insoluble Aβ were found not to correlate significantly with plaque burden as measured by histology potentially because most of the material stained by Campbell-Switzer silver staining is soluble in DEA (**Fig. 2K,M**).

### Aducanumab analog does not affect cerebral amyloid angiopathy in APP23 mice

We also investigated whether there was any effect of treatment with SUS or Adu or the combination on cerebral amyloid angiopathy (CAA). APP23 mice exhibit amyloid deposition on the vasculature with advanced age [31]. We found that there was no effect of Adu, or a combination of both on the number of blood vessels that were Campbell-Switzer positive and mice in all groups had significant deposition of amyloid on blood vessels (**Fig. 3A**). An average of twenty vessels were Campbell Switzer positive per section, with an average of 0.75% of the total area of the cortex taken up by amyloid-laden blood vessels and this did not differ between the groups (**Fig. 3B**). The findings for Adu not reducing CAA are in line with those data reported by Sevigny et al [5].

**Fig. 3.**
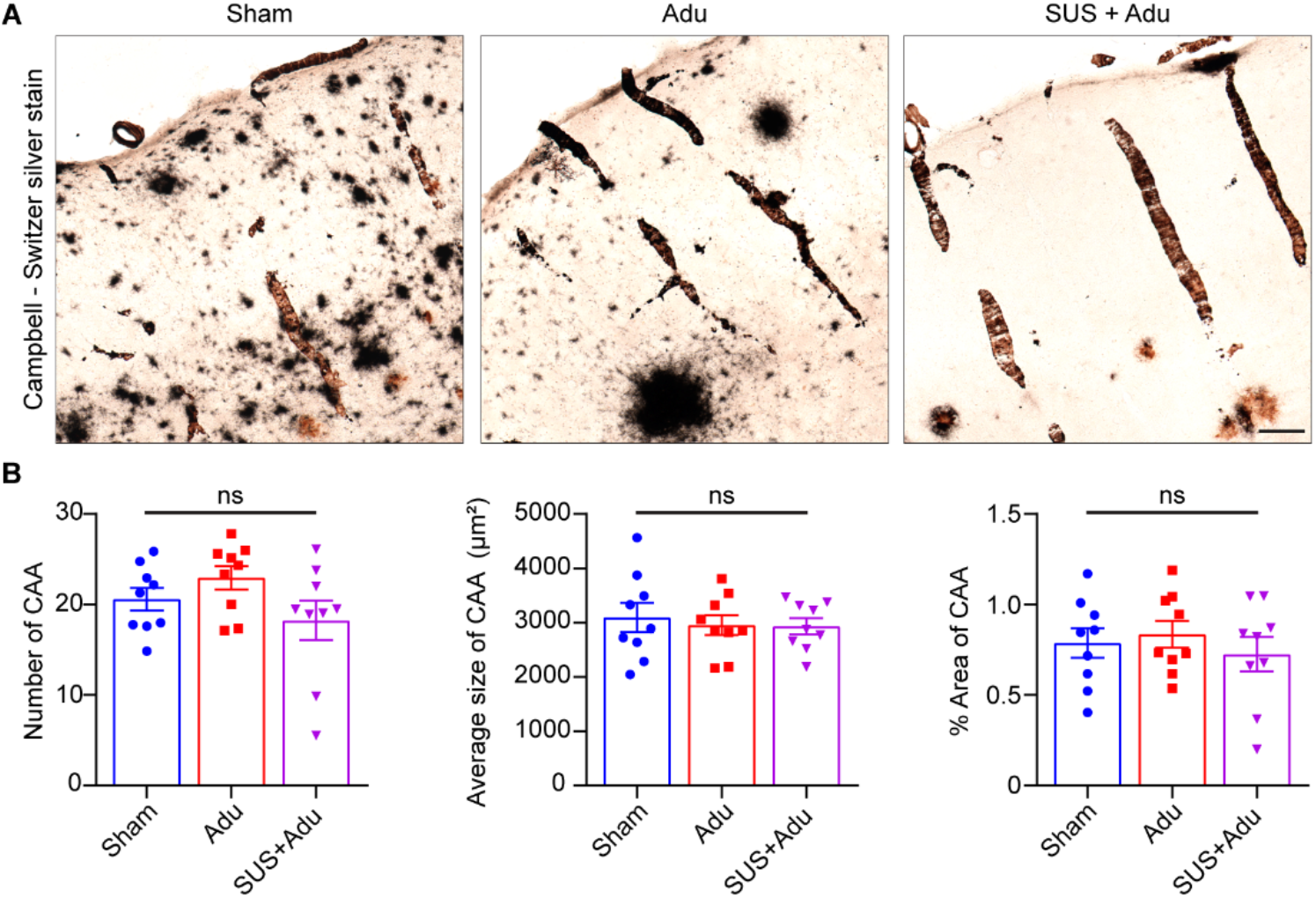
Aducanumab analog does not affect levels of cerebral amyloid angiopathy (CAA) in APP23 mice. (**A**) Representative Campbell-Switzer silver staining shows CAA in the cortex identified by a rod-like appearance, as well as meningeal CAA identified as open circles on top of the cortex. (**B**) Adu, whether administered with or without SUS, had no effect on the CAA number, average size, or percent area occupied by CAA. Data is represented as mean ±SEM. Statistical differences: *p<0.05. Data were analyzed with a one-way ANOVA and t-test.

### SUS markedly increases the amount of the Aducanumab analog in the brain

We also sought to determine the extent to which SUS was able to increase the amount of Adu in the brain. We therefore labeled Adu with Alexa Fluor 647 and injected 2.5 mg/kg fluorescently labeled Adu and 2.5 mg/kg unlabeled Adu at the last treatment session. In mice treated with Adu alone, fluorescently labeled Adu was faintly detectable by fluorescence microscopy and mainly confined to the outside edge of plaques (**Fig. 4A)**. In contrast, in SUS+Adu mice, fluorescently labeled Adu decorated the entirety of plaques and was easily detectable (**Fig. 4B)**. We also analyzed a subset of 5 mice per group to determine the area of the cortex that was positive for fluorescent Adu, revealing that 0.36% of the cortex in Adu treated mice was positive compared to 1.59% of the brain in SUS+Adu mice (p=0.0096, t-test). We also detected fluorescently labeled Adu in mice injected with Adu and SUS+Adu in the cortical lysate and found that levels were 4.32 ng/ml on average in the Adu group compared to 21.77 ng/ml in the SUS+Adu group (p=0.0175, t-test) (**Fig. 4C)**.

**Fig. 4.**
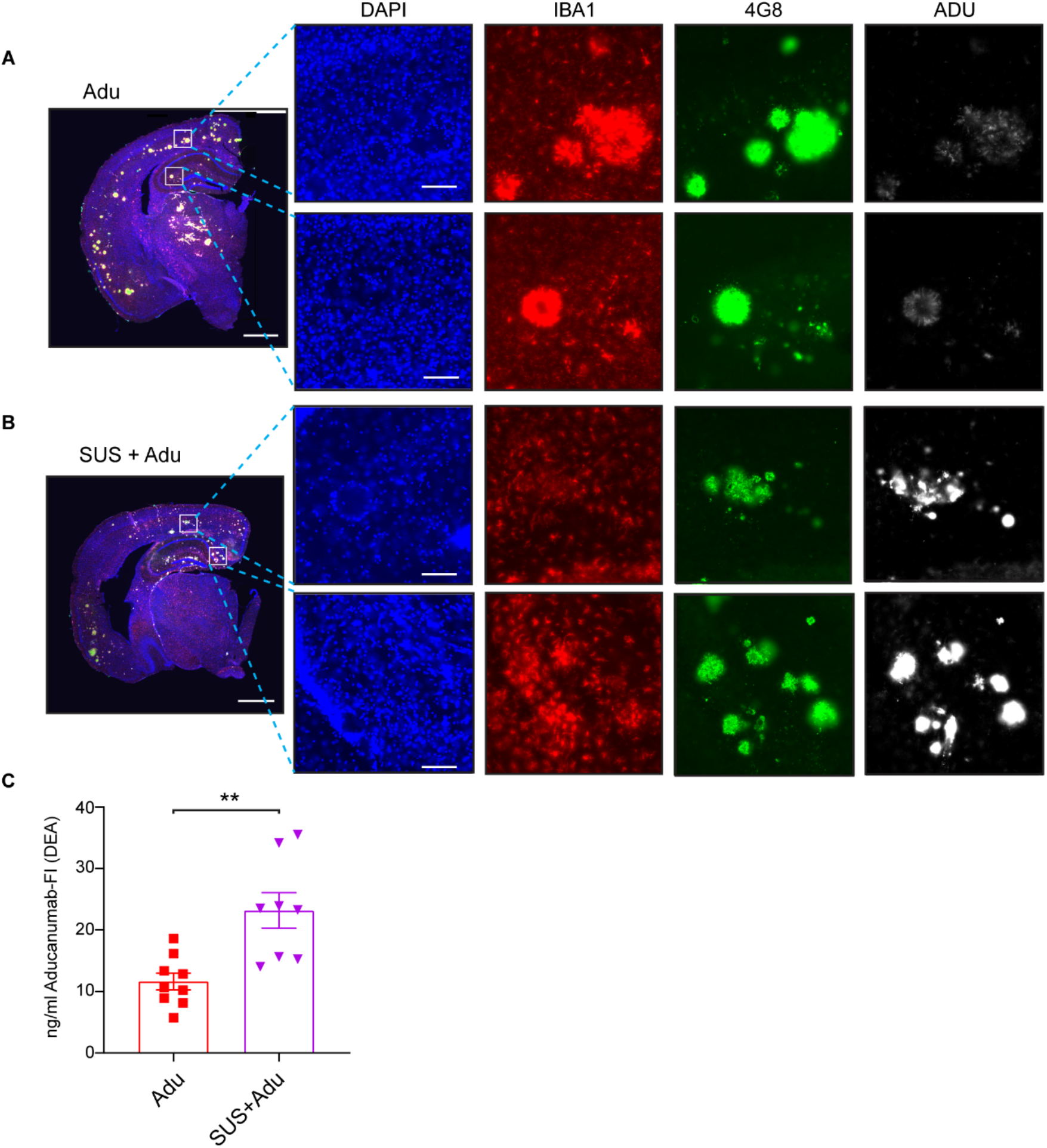
Scanning ultrasound (SUS) increases levels of the Aducanumab analog in the brain. (**A**) Fluorescently labeled Aducanumab analog is detectable in the brain when viewed in the entire brain (scale bars 1mm) and when visualized at higher magnification in the cortex and hippocampus (scale bars 100 μm). In APP23 mice treated with Adu alone the fluorescent Adu is detected bound to plaques, which were immunolabeled with 4G8 antibody. (**B**) Levels of Adu are higher when Adu was delivered with SUS in SUS+Adu-treated mice. The plaques of SUS+Adu-treated mice are decorated all over with Adu, whereas with Adu alone the Adu is mainly confined to the outsides of the plaque. Microglia as identified by IBA1 immunostaining are near plaques which have Adu bound to them. (**C**) Levels of fluorescent antibody in cortical brain lysate is increased in the SUS+Adu group compared to the Adu group. Data is represented as mean ±SEM. Statistical differences: *p<0.05. Data were analyzed with t-test. Adu N=9, SUS+Adu N=9.

## Discussion

Immunization strategies in AD have a long history, whereas therapeutic ultrasound has only recently been explored as a treatment modality. Our earlier work and that of others have revealed that ultrasound in combination with microbubbles, but in the absence of a therapeutic agent, can clear protein aggregates such as the hallmark lesions of AD, Aβ plaques [17, 18, 32, 33] and Tau neurofibrillary tangles [20, 22, 34]. We have also previously shown that the application of SUS achieves BBB opening throughout the brain [17] and, consequently, results in higher brain concentrations of antibodies in an IgG format, with up to 2% of the injected dose reaching the brain as compared to 0.2% without SUS [23]. Studies by us in Tau transgenic mice [22, 23] and work by others [24, 35] have suggested that ultrasound can also be used as an effective drug delivery tool to increase levels on antibodies in the AD brain.

Here, we used a multi-arm study, in which we compared the effects of SUS, an Aducanumab analog, Adu, delivered peripherally, and Adu delivered to the brain using SUS in APP23 mice with plaque pathology, using a sham treatment and wild-type mice as controls. In our study, SUS treatment had comparable effects as Adu treatment on plaque burden and behavior. We further found that in our treatment paradigm (nine treatments from age 13 to 22 months of age), when Adu was delivered across the BBB with SUS it more markedly reduced the amyloid plaque burden in the cortex of 22 month-old APP23 mice compared to the effects of either the antibody or SUS alone, concomitant with improvements in memory. For the first time we report data on the effect of an Aducanumab analog in a spatial memory task in plaque-bearing AD model mice. In our experiment we first performed the APA test of spatial memory and learning in 13 month-old APP23 mice to obtain a baseline, and then divided the mice into treatment groups based on their performance on the test which we reasoned would allow us to achieve greater power to detect improvements caused by the treatment due to reduced variability between the groups. We treated APP23 mice four times once per week and then repeated the APA test in which mice had to learn new spatial cues to avoid the shock zone. This experimental design allowed us to detect an improvement in mice treated with the combination of SUS and Aducanumab compared to sham-treated mice, and to detect improved performance in mice treated with combination compared to mice treated with Aducanumab without SUS, as well as to compare the effects of SUS to Aducanumab.

We were also interested in the effect of treatment on amyloid plaque burden, specifically at an advanced age in mice when plaque burden is more similar to that of an early AD patient, and therefore we aged the mice until 22 months of age. To conserve antibody, we performed five treatments over this period. Interestingly, plaque reduction in the hippocampus could be achieved with any of the three treatments (SUS, Adu and the combination), possibly reflecting the lower degree and later appearance of pathology in this brain area and suggests the plaque-lowering effect of SUS and Aducanumab are comparable in this brain area. Spatial memory was improved in 14 month-old APP23 mice following four weekly treatments with a combination of Adu and SUS. We have previously shown that a combination therapy using SUS and an anti-tau antibody increases the uptake of that antibody in IgG format 19-fold when measured one hour after treatment [23]. In line with these findings, we observed a five-fold increased levels of Adu in this study when administered in combination with SUS, which was measured three days after the treatment. The total increased uptake would likely be higher if measured at earlier time points post-injection, as IgG is cleared from the CNS or taken up by microglia as time progresses.

Delivering higher amounts of an anti-Aβ antibody into the brain should increase the efficacy of immunotherapy. This could in principle be achieved by strategies that bypass the BBB such as direct injection into the CNS. In one study, intracranial delivery of Aducanumab by topical application to the cortex was found to rapidly clear plaques in 22 month-old Tg2576 mice, whereas peripheral injections of Aducanumab at 10mg/kg repeated weekly from the ages of 18-24 months proved ineffective at clearing plaques, although they did restore physiological levels of intraneuronal calcium [36]. In contrast, plaque reduction was reported by Sevigny and colleagues when Aducanumab was delivered peripherally in Tg2576 mice. Plaques were reduced by 50% by weekly 10 mg/kg injections by treatment that began at 9.5 months of age and plaque burden was assessed at 15.5 months, suggesting that Aducanumab treatment may be less effective at removing plaques in mice with a substantial plaque burden compared to preventing plaque formation [5]. Our results show that administering a much lower cumulative dose of an Aducanumab analog than these authors had used is ineffective at clearing plaques in the cortex of APP23 mice when treatment is commenced at 13 months of age; however, plaques in the hippocampus which develop at a more advanced age were reduced. In contrast to peripheral injections alone, delivery of the Aducanumab analog using SUS led to a reduction in both cortical and hippocampal plaques, concomitant with increased brain levels of the antibody.

Efforts are currently underway in several laboratories to develop therapeutic ultrasound into a treatment modality for AD and other brain diseases, with ongoing clinical trials using an FDA-approved focused ultrasound system (ExAblate Neuro, Insightec) [16, 37] and implanted transducers [38] (Sonocloud, Carthera). These studies are applying ultrasound with microbubbles, but without a therapeutic agent such as an anti-Aβ antibody, with safe and effective BBB opening being used as primary endpoints, and Aβ clearance as a secondary endpoint. There are clearly several obstacles ahead such as being able to open a large enough brain area repeatedly and safely [8, 39]. Use of therapeutic ultrasound to open the BBB offers the possibility of achieving better brain uptake of a drug that has shown evidence of clinical efficacy, such as Aducanumab [5]. There is also the possibility of reducing the level of antibody administered facilitated by the use of ultrasound to achieve the same therapeutic outcome. This study provides further evidence also for the use of ultrasound opening of the BBB without a therapeutic agent by showing it is not inferior to anti-Aβ immunotherapy as both may act via increasing microglia phagocytosis of Aβ [5, 17]. More importantly, we believe that therapeutic ultrasound offers the prospect of combining therapeutic agents (such as an anti-Tau antibody and an anti-Aβ antibody) at levels below a safety threshold. What has not been discussed here is the potential of applying therapeutic ultrasound to target the brain in either a global or a more focused manner. We anticipate that once therapeutic ultrasound such as SUS has overcome the critical approval hurdles it may develop into a highly versatile and effective treatment therapy not only for AD but also for other brain diseases.

## Conclusions

An effective therapy for AD would reduce amyloid load and improve cognition. We show that Aducanumab and SUS alone have comparable ability to reduce amyloid levels. This presents SUS as a suitable treatment option for AD. The combination of SUS and aducanumab also improves cognitive function as measured by the APA test suggesting a combination trial using Aducanumab together with ultrasound. This may lead to increased brain levels of Aducanumab. Moreover, the two approaches may engage different (albeit shared) clearance mechanisms.

## Abbreviations

AD: Alzheimer’s disease
SUS: scanning ultrasound
Aβ: amyloid beta
ELISA: enzyme linked immunosorbent assay
Iba1: Ionised calcium binding adaptor molecule 1
IgG: immunoglobulin
Adu: Aducanumab mouse IgG2a chimeric analogue
Tg: Transgenic
W: wild-type

## Acknowledgments

We thank Linda Cumner for monitoring the animal Colony, and Rowan Tweedale for critically reading the manuscript.

## Author contributions

J.G, and G.L designed the research. G.L and W.K.K performed experiments. J.G, G.L, and W.K.K analyzed the data and wrote the paper.

## Availability of data and materials

The authors will make data available upon reasonable request.

## Ethics approval and consent to participate

All animal experimentation was approved by the Animal Ethics Committee of the University of Queensland (approval number QBI/554/17).

## Consent for publication

Not applicable.

## Supplementary information

**Figure S1.**
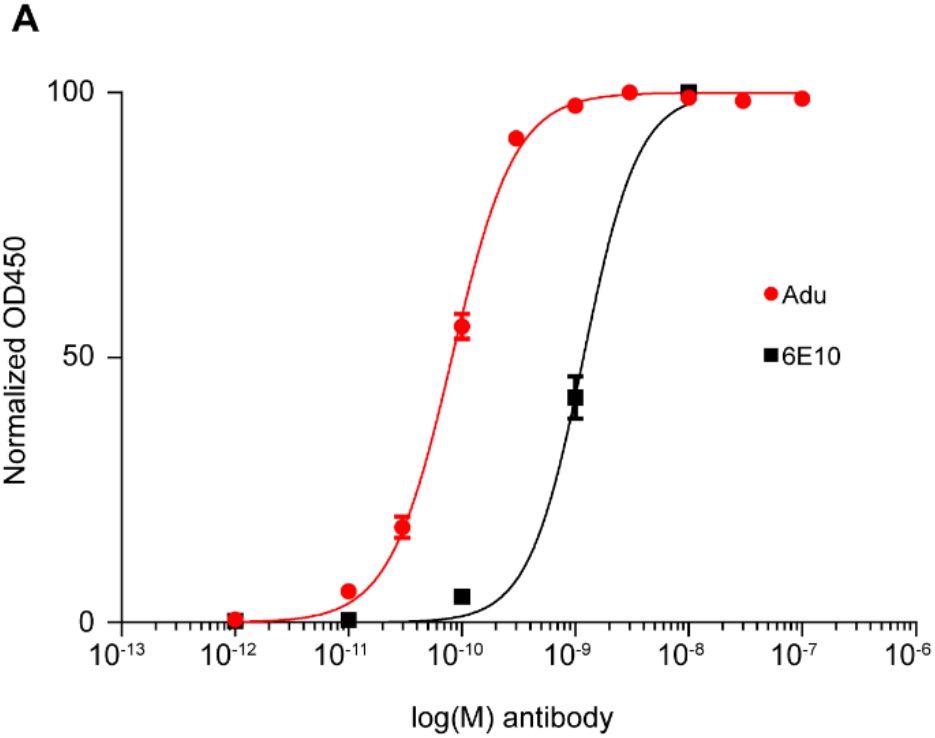
Affinity measurement of Aducanumab analog. The affinity of Aducanumab analog for fibrillar amyloid-β42 was measured by ELISA and compared to the antibody 6E10.

**Figure S2.**
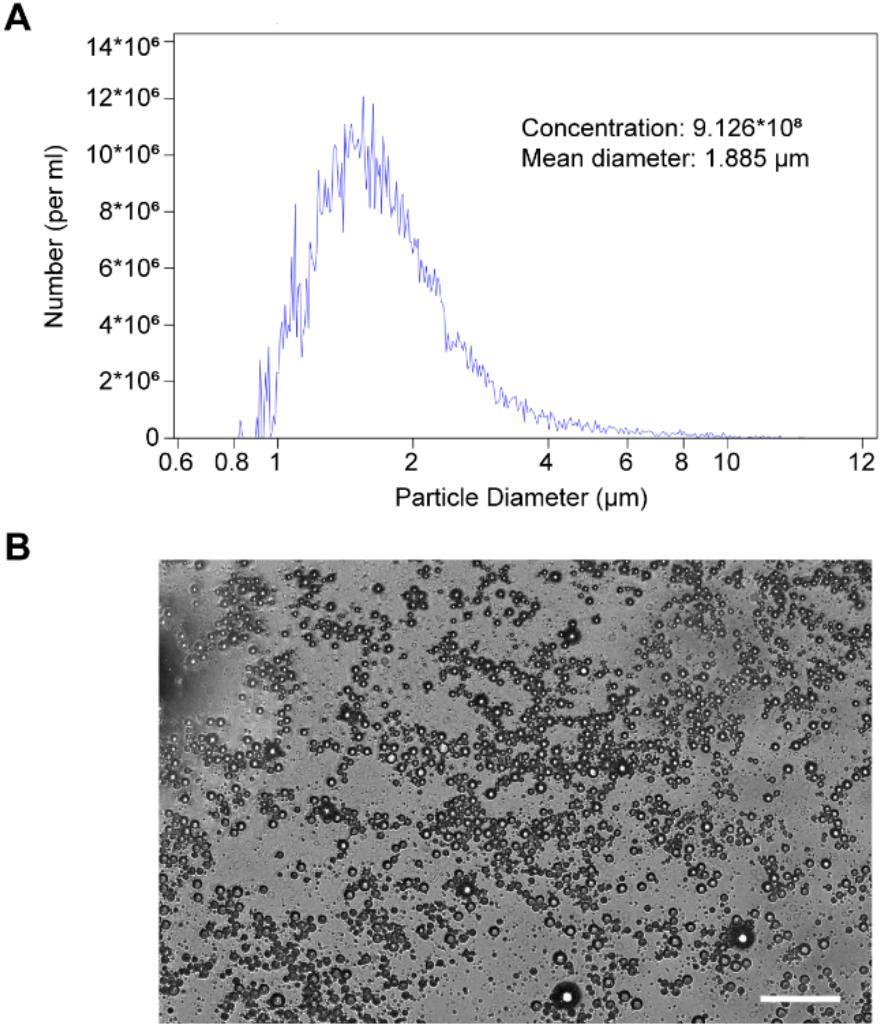
Characterization of microbubbles. In-house prepared microbubbles were analyzed by Coulter Counter.

